# The use of mixture-density networks in the emulation of complex epidemiological individual-based models

**DOI:** 10.1101/551440

**Authors:** Christopher N Davis, T Deirdre Hollingsworth, Quentin Caudron, Michael A Irvine

## Abstract

Complex, highly computational, individual-based models are abundant in epidemiology. For epidemics such as macro-parasitic diseases, detailed modelling of human behaviour and pathogen life-cycle are required in order to produce accurate results. This can often lead to models that are computationally-expensive to analyse and perform model fitting, and often require many simulation runs in order to build up sufficient statistics. Emulation can provide a more computationally-efficient output of the individual-based model, by approximating it using a statistical model. Previous work has used Gaussian processes in order to achieve this, but these can not deal with multi-modal, heavy-tailed, or discrete distributions. Here, we introduce the concept of a mixture density network (MDN) in its application in the emulation of epidemiological models. MDNs incorporate both a mixture model and a neural network to provide a flexible tool for emulating a variety of models and outputs. We develop an MDN emulation methodology and demonstrate its use on a number of simple models incorporating both normal, gamma and beta distribution outputs. We then explore its use on the stochastic SIR model to predict the final size distribution and infection dynamics. MDNs have the potential to faithfully reproduce multiple outputs of an individual-based model and allow for rapid analysis from a range of users. As such, an open-access library of the method has been released alongside this manuscript.

**Author summary:** Infectious disease modellers have a growing need to expose their models to a variety of stakeholders in interactive, engaging ways that allow them to explore different scenarios. This approach can come with a considerable computational cost that motivates providing a simpler representation of the complex model. We propose the use of mixture density networks as a solution to this problem. These are highly flexible, deep neural network-based models that can emulate a variety of data, including counts and over-dispersion. We explore their use firstly through emulating a negative-binomial distribution, which arises in many places in ecology and parasite epidemiology. We then explore the approach using a stochastic SIR model. We also provide an accompanying Python library with code for all examples given in the manuscript. We believe that the use of emulation will provide a method to package an infectious disease model such that it can be disseminated to the widest audience possible.

## Introduction

Complex individual-based models abound in epidemiology. These can often include a mixture of different distributions leading to a high-dimensional output space with possibly multi-modal distributions. As an example of this, consider the relatively simple stochastic Susceptible-Infected-Recovered (SIR) model [1]. This is where an epidemic occurs where each individual has a random chance of becoming infected and further contributing towards the ongoing epidemic. Depending on initial conditions and parameters, an epidemic may be observed or the initial number of infected individuals recover before a large-scale epidemic can occur (stochastic fade-out) [2]. This example would then lead to a multi-modal total number of possible infected individuals to account for samples where an epidemic did not occur and when it did.

Other examples of complex, individual-based models occur in macro-parasitic diseases. These diseases will often have highly heterogeneous parasite distributions amongst its hosts, which therefore requires explicit individual hosts to be modelled in order to directly understand the consequences and implications of these distributions [3]. Compounding this, is that many of these diseases are controlled through mass drug administration, where coverage, adherence and demographic factors can play a role in the outcome of a program [4,5].

Finally, many examples of individual-based models exist in sexually-transmitted infectious disease, such as HIV, HPV, gonorrhea, and syphilis [6]. Here complexities such as heterogeneity in risk, partnership-formation, sexual contact networks, sero-sorting behaviour, and heterogeneity in interventions can lead to dramatically different outcomes that require modelling.

Coupled with this increasing number of computationally-expensive models is the move towards models being more open to non-experts in order to allow exploration of key concepts and outputs of a model. There has been an increasing call for more models to be outward facing to be used by policy makers and other non-modellers [7]. However, despite this there remains significant technological barriers to be able to perform this in general, often requiring skill in multiple programming languages and software development [8]. In particular, one of the technological barriers is the speed at which model simulations of a given scenario can occur. This introduces the idea of using emulation in lieu of model computation [9–13]. This is where the individual-based model is replaced by a statistical model that is more computationally efficient to simulate from. Training of these statistical models can be difficult and will often rely on unimodal or normality assumptions on the outcome distributions of the model [14].

As individual-based models become more complex, the necessary computational costs increase. This can often lead to only a small number of scenarios being explored with relatively few replicates used to estimate uncertainty within the model. This is particularly challenging when many simulation runs need to be performed such as in an inference scheme like approximate Bayesian computation [15]. Computational speed-up can be performed by making certain approximations within the model, such as taking the deterministic limit for a process that has relatively large numbers, or again through the use of emulation [4,14].

The main concept of an emulator is to fit a statistical regression model to the inputs and outputs of the individual-based model and then evaluate the computationally much faster model instead [16]. For example, an individual-based model may have *m* number of inputs (*x*_0_, …, *x*_*m*-1_), with a corresponding single output *y*. We may also assume that for a given set of inputs the output is normally-distributed. We may then emulate the model using the following linear regression

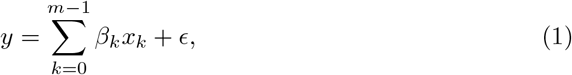

where *ϵ* ∼ *N* (0, *σ*^2^). For all but simple individual-based models the linearity assumption may be too restrictive. The fixed *β*_*k*_*x*_*k*_ terms may be replaced with a Gaussian process (GP), where values are allowed to vary across the input space, with the assumption that the closer input points are together, the more correlated will be the outputs. GPs have had a number of successes within emulation of epidemiological models [14,16–19].

A GP has a certain number of disadvantages, however. They first assume that the outputs of a model for a given set of inputs are normal. This would not be able to take into account the multi-modality or heavy-tailedness of certain data. There may also be some restrictive assumptions on the smoothness of the correlation between two points in input space. Here we propose the use of a mixture density network to overcome some of these issues. The idea explored in this study is to replace the linear regression component with a neural network that is flexible enough to capture complex relationships and replace the simple normal distribution with a mixture of distributions that provides a more general family of distributions for the model output [20]. A comparison of GPs and mixture density networks is shown in Table 1.

**Table 1.** A comparison of mixture density networks and Gaussian processes.

| Mixture density network | Gaussian process |
| --- | --- |
| Can emulate multi-modal distributions. | Only suitable for uni-modal distributions. |
| Flexible output distribution mixture allows for application to different data types, such as overly-dispersed, finite domain or discrete. | Output distribution is normal. |
| Require large training set to capture output. | Suitable for small data sets. |
| Good scaling properties. | Scales poorly with data size. |
| Hyperparameters need to be tuned in training process. | Hyperparameters need to be tuned in training process. |
| The training process is stochastic. | For given parameters, the method is optimised exactly. |
| Uncertainty hard to quantify. | Directly measures uncertainty. |

The manuscript is segmented as follows: We first introduce the concept of a mixture-density network and how it relates to an individual-based model, we then apply this to a number of simple examples to demonstrate its use. We then apply the model to a stochastic SIR model and emulate the final size distribution. Finally we demonstrate its use on estimating the distribution of susceptible and infected individuals in a model with vaccination. All analyses presented within the manuscript were conducted within the package framework and example code is given (see supporting information S2).

## 1 Methods

### Introduction to mixture density networks

Mixture density networks are built from two components: a neural network and a mixture model. We begin by introducing the concept of a mixture model, these are a model of probability distributions built up from a weighted sum of more simple distributions. More concretely consider a one-dimensional distribution with *m* mixture components. The probability density function (pdf) *p*(*x*) is represented by the *m* pdf indexed by *j p*_*j*_(*x*), with weights Π = {*π*_0_, …, *π*_*m*-1_} by the following equation

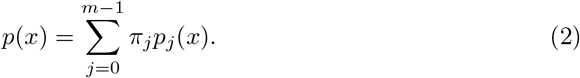

Typically these probability distributions will be parameterised by a series of parameters that reflect the shape and location of the distribution Θ = {*θ*_0_, …, *θ*_*m*-1_}. The full parameterised model may therefore be written as

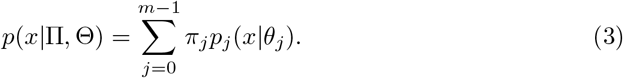

As an example, each *p*_*j*_ could be a normal distribution parameterised by a mean *µ*_*j*_ and a variance *σ*_*j*_. The mixture model would then have the following form,

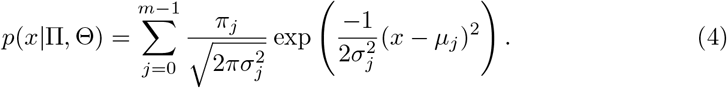

In general these mixture distributions are multi-modal and can be fitted directly to some data **x** = (*x*_0_, …, *x*_*n*-1_). The corresponding likelihood is calculated as

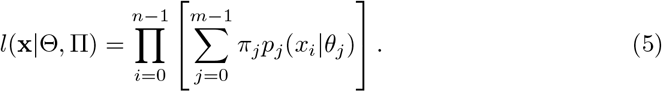

Fitting can then typically proceed using expectation-maximization [20]. For our purposes, we have an individual-based model *M*, with some input *α*, that produces stochastic realisations *y* ∼ *M* (*α*). We therefore wish to derive a relationship between the input parameters *α*, and the mixture density weights *π*_*j*_(*α*) and density parameterisations *θ*_*j*_(*α*). This could potentially be done with a separate regression for each of the density parameters and weights, however this would fail to capture the corresponding relationships that would exist between each parameter and weight. We can therefore model these using a neural network which is able to provide flexible fitting for arbitrarily complex relationships by the universal approximation theorem [21]. A mixture density network is therefore defined as a mixture model, where the mixture components are modelled using a neural network.

Fig. 1 provides an overview of the mixture density network construction. The inputs of the model *α* are initially fed into the mixture density network (three such inputs in the example diagram). These are then passed through a number of hidden layers in the neural network, which provide a compact representation of the relationship between the inputs and the unnormalized inputs into the mixture model. These distribution parameters are then passed through a normalisation layer, where the weights of the mixture are transformed such that they sum to one and the shape parameters are transformed so that they are positive. These parameters are then used to construct the mixture model, where samples can be drawn from or statistics such as mean and variance can be calculated for a given input. For multiple outputs the final layer can be copied with independent parameters for the number of outputs being considered. Note that a number of aspects of the mixture density network need to be specified including the number of input parameters, the dimension of the output, the distributions used in the mixture density, and the number and size of the hidden layers.

**Fig 1.**
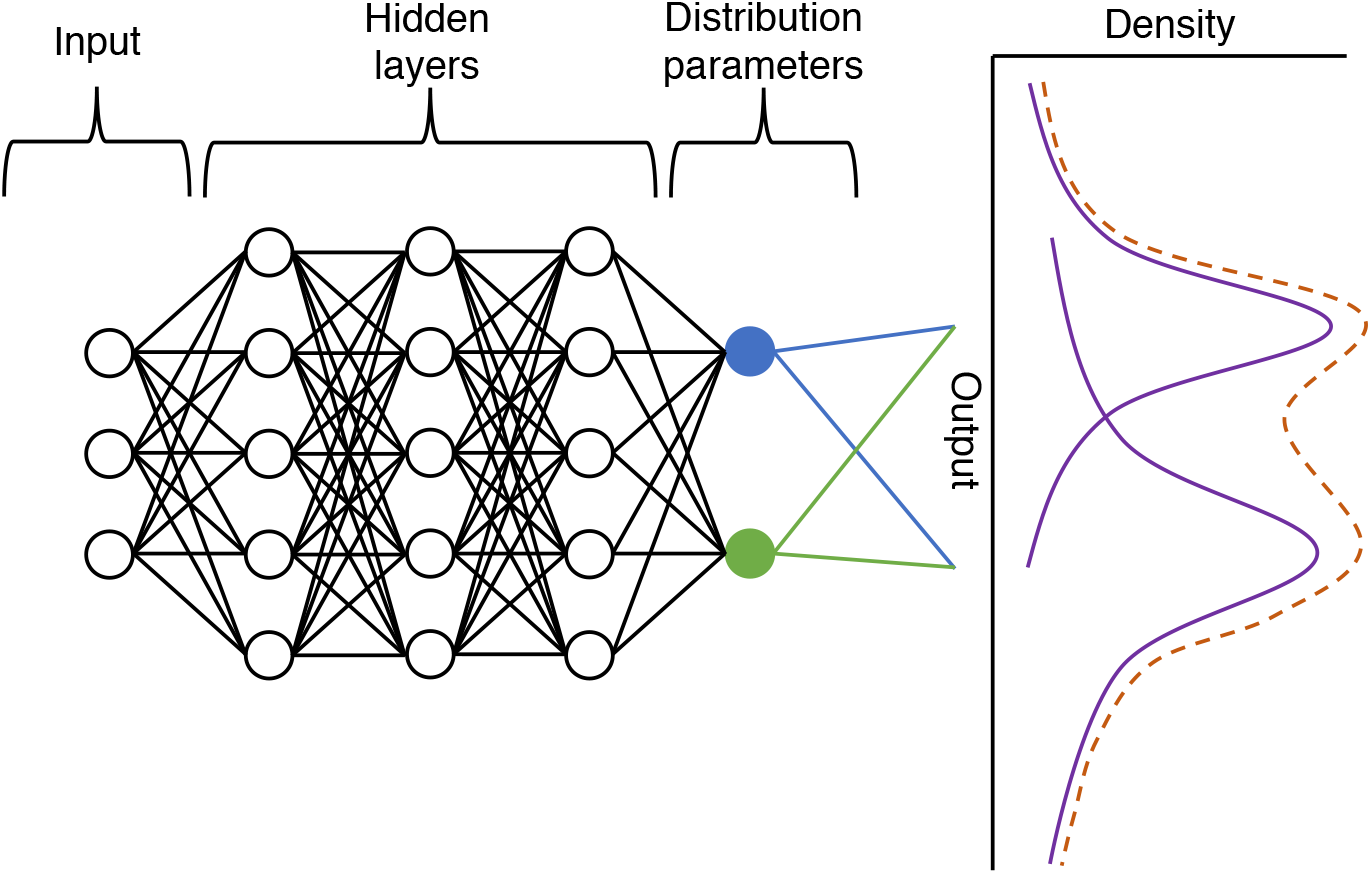
Mixture density network that emulates a model with three inputs and a one-dimensional output with two mixtures. The inputs are passed through two hidden layers, which are then passed on to the normalized neurons, which represent the parameters of a distribution and its weights e.g. the mean (shown in blue) and variance (shown in green) of a normal distribution. These parameters are then used to construct a mixture of distributions (represented as a dashed line).

The MDN can then be fit to the following objective loss function, which is equivalent to maximising the likelihood given in Eq. 5,

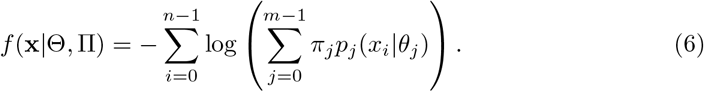

Note that provided *p*_*j*_ is differentiable with respect to *θ*_*j*_, this loss represents a differentiable function. Standard techniques based on stochastic gradient descent can then be applied in order to optimize the weights of the network with respect to this loss [22].

### Performance on a simple model

In order to examine how a fitted mixture-density network can capture the broad statistical properties of a distribution, where the underlying mixture distributions differ significantly from the true distribution, we explored fitting to a negative-binomial model. The negative-binomial can be parameterised by a mean *m* and a shape parameter *k* using the following probability mass function,

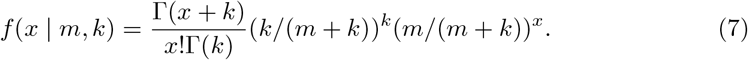

~~~
import pydra #import MDN emulator library
x = input data () # import input data with shape (# data points, # inputs)
y = ouput data () # import output data with shape (# data points,)
’ ’ ’
construct an emulator with 20 mixtures, three layers with 64 neurons and
    two inputs with one output that has a gamma mixture distribution
’ ’ ’
model = pydra.Pydra (cluster_size =20, output_size=1,layers = 3,input_size = 2,
                           dense_layer_size=64,output_distributions = [’Gamma’],
    print_summary=True)

# fit for 150 epochs with batch size 50
model.fit(data, y, epochs=150, batchsize=50, verbose=1)
~~~

**Listing 1.** Code for negative binomial distribution example

The parameter *m* defines the mean of the distribution and the shape parameter *k* controls the heterogeneity of the distribution, where the variance is *m*(1 + *m/k*). As *k* goes to infinity, the distribution approaches the Poisson distribution.

A mixture density network was fitted to the negative-binomial distribution in the following way. A mixture density network with 20 Gamma mixtures with 3 dense layers of 64 neurons was constructed. Data were sampled as 1000 (*m, k*) pairs uniformly at random from *m* with range 0–100 and from *k* with range 0.01–5. Fitting was performed for 150 epochs with batch-size 50.

In order to compare the statistical properties between the true negative-binomial distribution and the mixture density network emulator a number of tests were devised. First the mean and variance of each distribution were compared by fixing one of the parameters to the mid-point of the parameter range and varying the other parameter. In order to statistically compare between a sample generated from the true process and generated from an emulator, the two-sided Kolmogorov-Smirnov test was performed on two samples of size 100 across a range of input values for 100 replicates [23]. The true cumulative density function and the empirical cumulative density functions were also compared.

This experiment broadly captures how well the MDN can emulate a distribution significantly different to its underlying mixture-distribution, as well as how capable it is to adequately deal with highly heterogeneous data, which can complicate model fitting [15]. Example code using the accompanying open-source library is given in Listing 1.

### Stochastic SIR model

The stochastic susceptible-infected-recovered (SIR) model is a standard epidemiological model for the infection dynamics of an epidemic. It is a Markov process where each individual of a population of fixed size *N* can be in any one of the 3 states – *S, I* or *R*. Due to the stochasticity of the model, the output is multi-modal, where identical input parameters can have both quantitatively and qualitatively different outputs, depending on whether an epidemic does or does not occur. The transitions for the process are defined by:

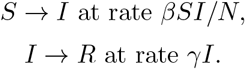

We generate multiple realisations of the stochastic SIR model and use this data to fit a series of MDNs such that we can evaluate the performance of the emulation model.

We note that while there are already fast methods to draw realisations of the process (Gillespie, tau-leap method), we assess this method to understand the benefits for when we want to emulate data from a more computational intensive model [24,25]. Here, we consider training 3 different MDNs for different inputs and output distributions.

Firstly, we consider the final size distribution of the epidemic model, given as the total number of individuals that have been infected (calculated as *N* (∞) – *S* (∞)). We take 10,000 realisations of the simulated process using model parameters *β*, sampled from *U* (0, 1), and *γ*, sampled from *U* (0.1, 1), with a population size of *N* = 1000. To fit a MDN for the final size given these inputs, we choose a mixture of 20 binomial distributions, where the binomial parameter *n* = 1000 is fixed and *p* is learnt from the MDN. We use binomial distributions as the final size is integer valued with a maximum value of the population size. We train on a network with 3 dense layers of 64 neurons for 150 epochs with a batch size of 50.

Due to the multi-modality, unlike for the negative binomial distribution, calculating the mean and variance of the whole distribution is not a useful concept and so we consider the similarity between the simulated and emulated distributions, by sampling from both. To quantify their equivalence, the two-sided Kolmogorov-Smirnov test was performed on two samples of size 100 (for simulated and emulated data) for randomly sampled *β* and *γ* values for 100 replicates. Furthermore, we compare empirical CDFs and since the distribution is bi-modal, compare the proportion of realisations were the final size is greater than 10% of the population size, indicating that an epidemic has occurred. These tests were performed across a range of *R*_0_ = *β/γ* values.

Secondly, we explored the infection dynamics across time. We take 10,000 realisations of the simulated process using model parameters *β*, sampled from *U* (0, 1), *γ*, sampled from *U* (0.1, 1), and time *t* sampled as a random integer between 1 and 100. The population size remains fixed with *N* = 1000. A MDN was trained to have 2 outputs, the prevalence of susceptible and infected individuals, for the 3 inputs of *β, γ* and time (scaled to be between 0.01 and 1). The prevalence of recovered individuals can be inferred from this given the fixed population size. Since the aim was to learn the prevalence rates, the number in each compartment divided by the population size, we use a mixture of 20 Beta distributions, as the Beta distribution is defined on the interval [0,1]. The MDN has 3 dense layers of 64 neurons and is trained for 50 epochs with a batch size of 50. The simulated and emulated distributions are compared and a 2 sample KS-test was performed on the output.

Finally, we added an additional transition for the process to model vaccination of susceptible individuals, which is given by

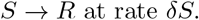

We then used the same method as before, with the 2 extra inputs of the vaccination rate *δ* sampled from *U* (0, 0.01) and population size sampled as a random integer between 1 and 1,000 (both linearly scaled to have a maximum of 1). Again, the simulation and emulation wass compared and a KS-test performed.

## Results

### 1.1 Simple model

The fitted gamma-mixture density network emulator was able to broadly capture the mean and variance of the distribution over a range of inputs parameters (Fig. 2a & 2b). There were some notable deviations to these statistics however, where the parameters were near the edge of the range. In particular, where *k <* 1, both the mean and variance begin to differ significantly from the true distribution where the variance increases to infinity as *k* goes to zero (Fig. 2b).

**Fig 2.**
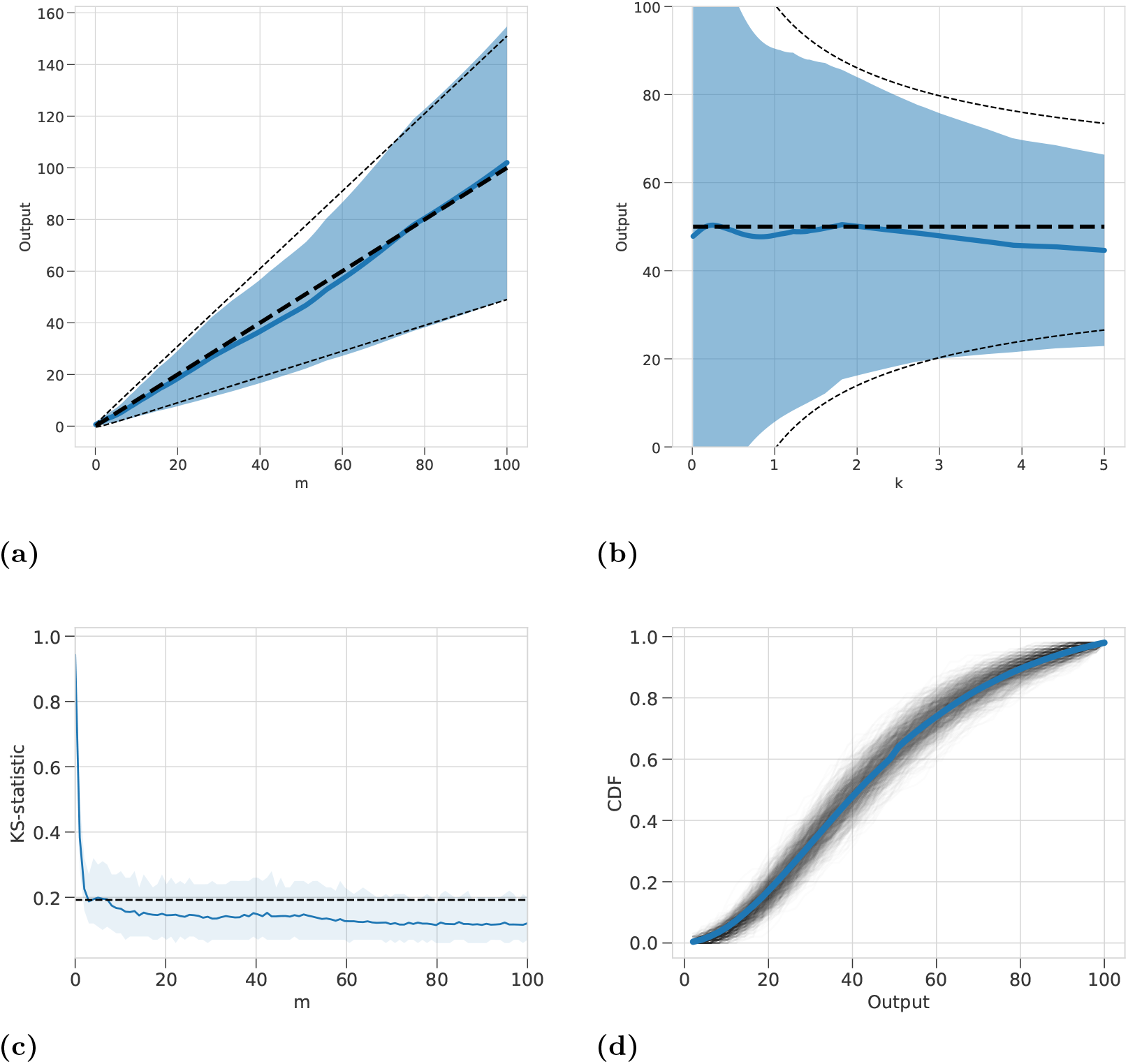
Gamma-mixture-density network output emulating a negative-binomial model. **(a)** For fixed shape parameter *k* = 2.5, the distribution of output from MDN is shown in blue (mean = solid line, variance = shaded region), the theoretical values are shown as a black dashed line (mean= bold line, variance = normal line) **(b)** For fixed mean parameter *m* = 50, the distribution of output from MDN over a range of *k* values is shown in blue (mean = solid line, variance = shaded region), the theoretical values are shown as a black dashed line (mean= bold line, variance = normal line) **(c)** Corresponding two-sample KS-statistic where sample of 100 points are drawn from a negative-binomial and the MDN over a range of *m* values. 100 replicates are used to estimate a mean KS-statistic and a 95% range. Dashed line represent significance at *α* = 0.05, with values less indicating the two-samples do not differ significantly. **(d)** Example empirical cumulative density functions drawn from 100 samples of MDN with inputs *m* = 50 and *k* = 2.5. 1000 empirical CDF are shown as black transparent lines and true CDF is shown as a blue solid line.

The KS-statistic is below the significance level over a broad range of *m* values indicating a sample drawn from the true process and the emulator are similar (Fig. 2c). Only when *m* is close to zero at the edge of the range of training data does the distribution differ significantly from the true distribution according to the KS-statistic. This can also be broadly shown by plotting example empirical CDF against the true CDF (Fig. 2d).

### 1.2 Stochastic SIR model

The three MDNs were capable of emulating the behaviour of the stochastic SIR model. Sampling from the output distribution of the final size of an epidemic, there was strong agreement with the results of sampling from the actual simulation across the full range of *R*_0_ values (Fig. 3a). This is corroborated by the results of the KS-test, where the KS-statistic lies below or close the the significance level that the samples could be drawn from the same distribution for different *R*_0_ values (Fig. 3b). The result was stronger for both small and large *R*_0_ values, with some divergence for the intermediate values, where the final size takes a larger range of values. Comparison of the proportion of samples that exhibit stochastic fade out as opposed to the emergence of an epidemic was determined where the final size reaches at least 10% of the population (Fig. 3c). These proportions matched closely, with the emulation only slightly deviating to a higher than expected proportion for large *R*_0_ values. The results also matched for the CDFs of the two distributions, shown for a fixed *R*_0_ = 2 with *β* = 0.4, *γ* = 0.2 (Fig. 3d).

**Fig 3.**
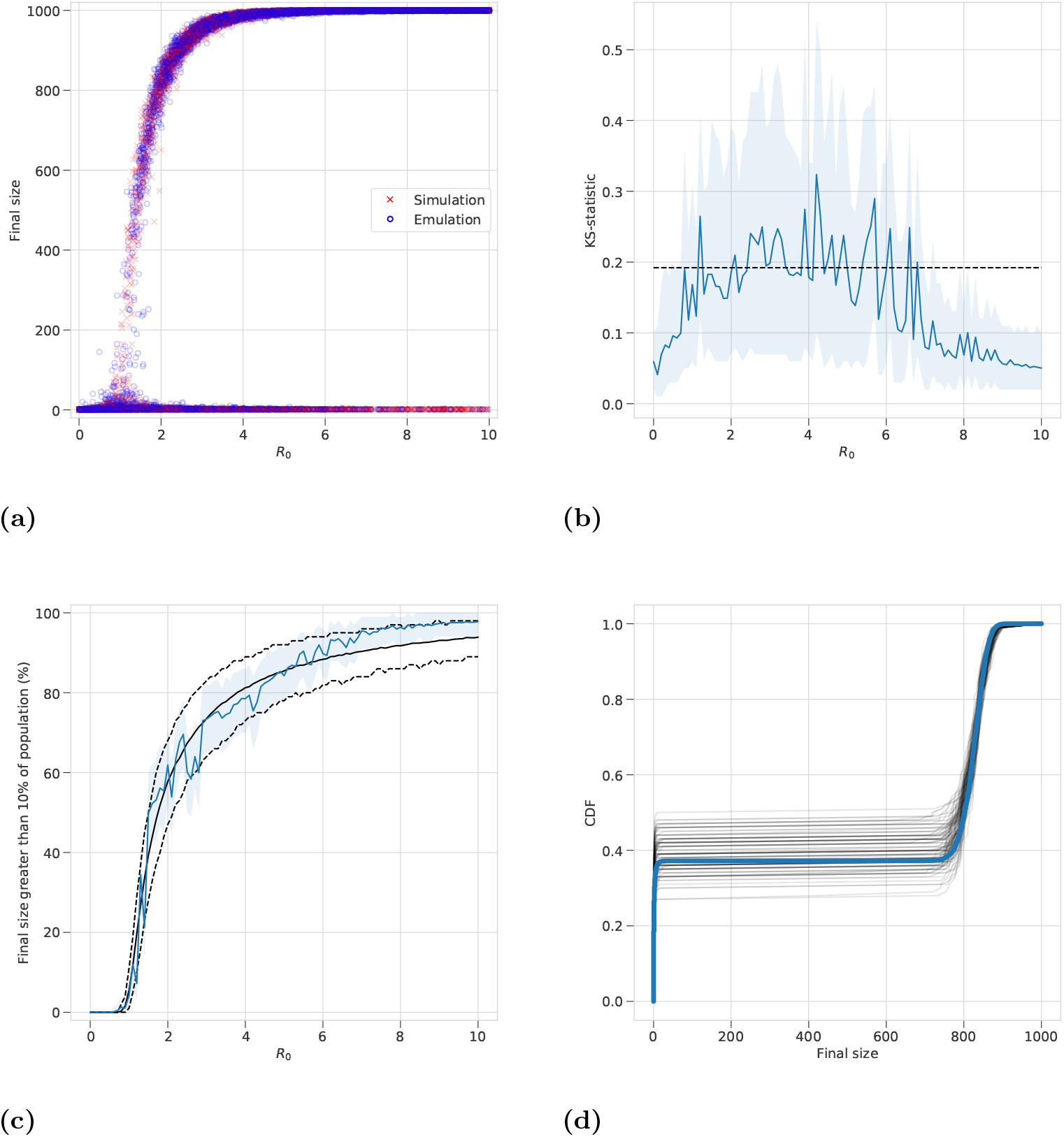
Binomial-mixture-density network output emulating the final size distribution of a stochastic SIR model. **(a)** For random uniform sampling over *β* and *γ* a sample of the output from MDN across values for the basic reproductive number *R*_0_ = *β/γ* are shown in blue and the directly simulated values are shown in red. **(b)** Corresponding two-sample KS-statistic where sample of 100 points are drawn from a negative-binomial and the MDN over a range of *R*_0_ values. 100 replicates are used to estimate a mean KS-statistic and a 95% range. Dashed line represent significance at *α* = 0.05, with values less indicating the two-samples do not differ significantly. **(c)** The percentage of 1,000 realisations of the stochastic SIR model with final size greater than 100 is shown in black with dashed line showing a 95% range. Emulated results are shown by the blue line with a 95% range. **(d)** Example empirical cumulative density functions drawn from 100 samples of MDN with inputs *β* = 0.4 and *γ* = 0.2. 1000 empirical CDF are shown as black transparent lines and true CDF is shown as a blue solid line.

The addition of emulating the actual infection dynamics against time shows that a single trained MDN captures the complexity of varying *R*_0_ by qualitatively reproducing when an epidemic occurs in the correct timescales (Figs. 4a – 4d). However, statistically comparing the two output distributions for both susceptible and infected individuals, the KS-statistic does increase above the significance level (Figs. 4e & 4f). This is in part as the beta distribution is a continuous distribution, whereas the simulated prevalence values are all fractions of the population size *N* = 1000. The effect of this is particularly stark for small time *t* values where the number of individuals will always be exactly *S*_0_ = 999 and *I*_0_ = 1, whereas there will be some variation outside these values in the emulation as it is not integer valued. Rounding the emulated values or choosing an integer-valued output distribution helped to improve the KS test results (see supplementary).

**Fig 4.**
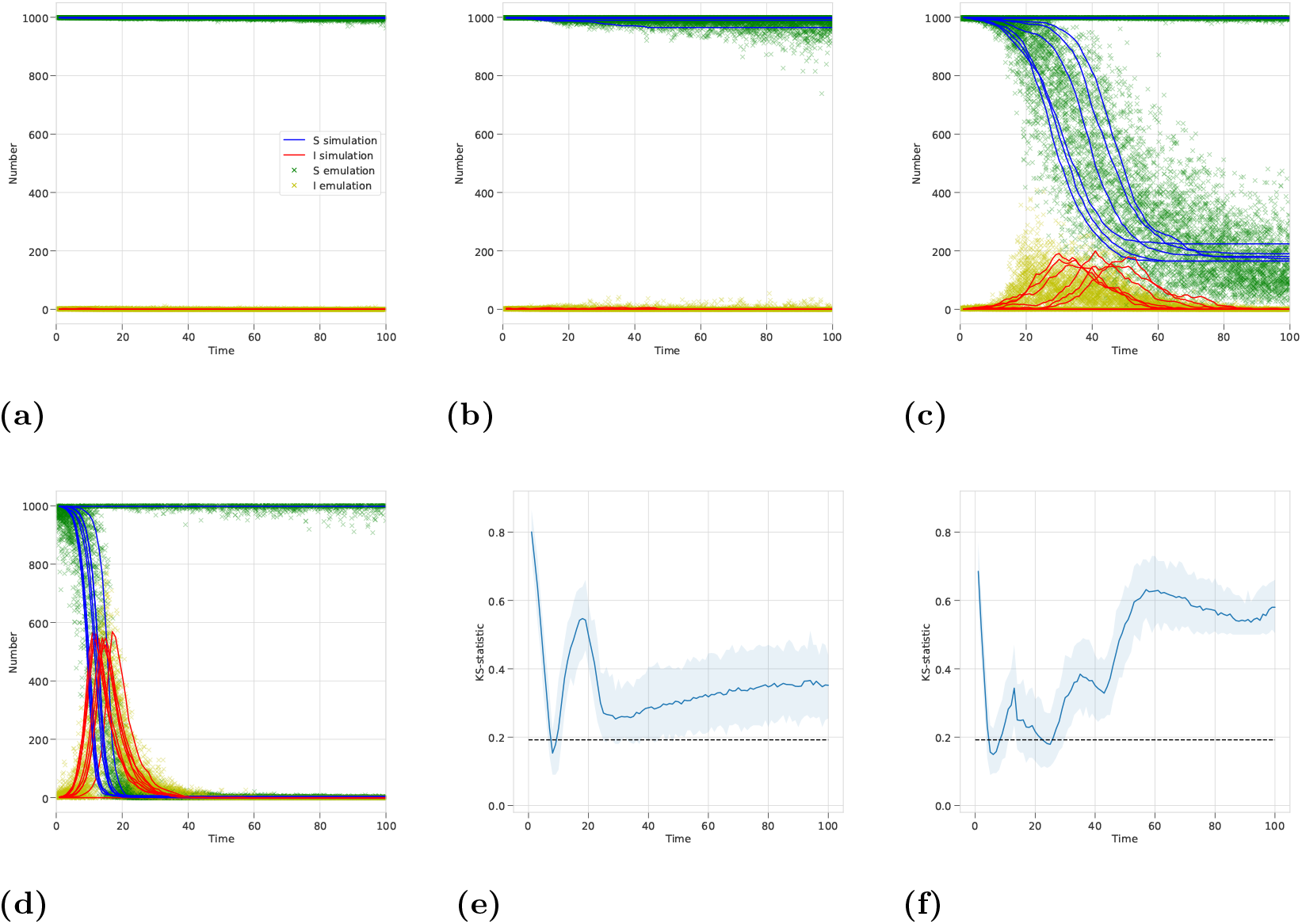
Beta-mixture-density network output emulating the infection dynamics with time for a stochastic SIR model. **(a-d)** A comparison of simulation results with sampled MDN output for fixed *γ* = 0.2 and *N* = 1000 and different *β* values such that **(a)** *R*_0_ = 0.5, **(b)** *R*_0_ = 1.0, **(c)** *R*_0_ = 2.0, **(d)** *R*_0_ = 5.0. **(e-f)** Two-sample KS-statistic where sample of 100 points are drawn from a negative-binomial and the MDN over a range of time *t* values. 100 replicates are used to estimate a mean KS-statistic and a 95% range. Dashed line represent significance at *α* = 0.05, with values less indicating the two-samples do not differ significantly. Tests are for **(e)** number of susceptible people and **(f)** number of infected people.

Furthermore, the emulation coped well with learning to reproduce the distributions with further model parameters, vaccination rate *δ* and population size *N*, since inputting different values for *β, γ, δ* and *N* into the trained MDN qualitatively reproduces the infection dynamics of the simulation across all time values *t* (Figs. 5a – 5d). Similarly to the MDN without variable vaccination and population size, the results of the KS test suffer from a similar problem of comparing discrete and continuous distributions, but the resulting distributions were very close at some values (Figs. 5e & 5f).

**Fig 5.**
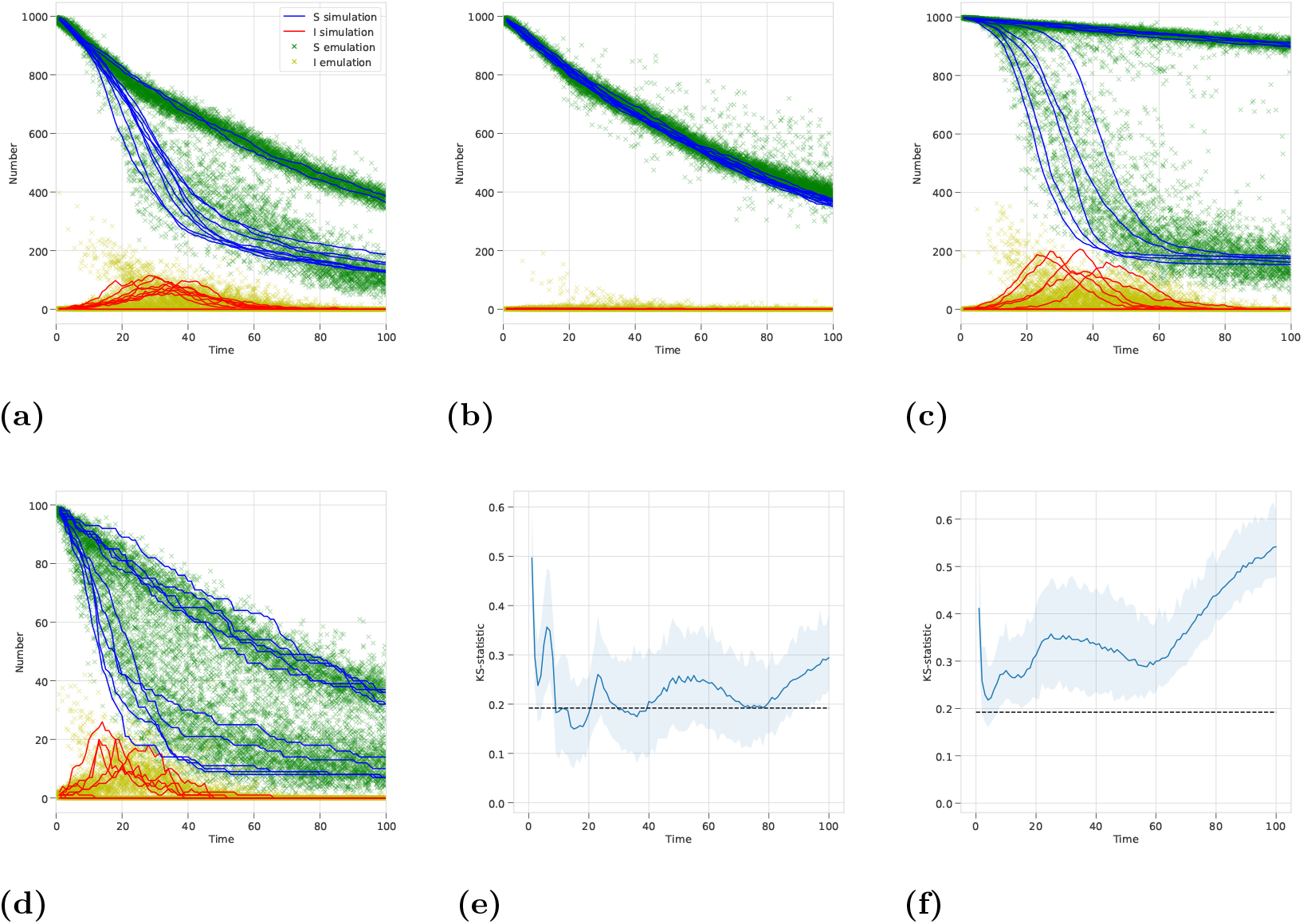
Beta-mixture-density network output emulating the infection dynamics with time for a stochastic SIR model. **(a-d)** A comparison of simulation results with sampled MDN output for fixed *γ* = 0.2 and different *β, δ* and *N* values such that **(a)** *R*_0_ = 2.0, *δ* = 0.01 and *N* = 1000, **(b)** *R*_0_ = 1.0, *δ* = 0.01 and *N* = 1000, **(c)** *R*_0_ = 2.0, *δ* = 0.001 and *N* = 1000, **(d)** *R*_0_ = 2.0, *δ* = 0.01 and *N* = 100,. **(e-f)** Two-sample KS-statistic where sample of 100 points are drawn from a negative-binomial and the MDN over a range of time *t* values. 100 replicates are used to estimate a mean KS-statistic and a 95% range. Dashed line represent significance at *α* = 0.05, with values less indicating the two-samples do not differ significantly. Tests are for **(e)** number of susceptible people and **(f)** number of infected people.

## Discussion

Model emulation is quickly becoming an important and necessary method within infectious disease epidemiology due to the increased use of complex, computationally-intensive models, increased use of direct data fitting requiring many model queries, and increased demand for models to be made directly available to knowledge users [26]. We have explored the use of mixture density networks (MDN) in order to provide a scalable, flexible solution to this type of emulation [27]. These are mixture models where the underlying parameters of the mixture are neural networks. This allows the significant progress in neural networks and deep learning to be incorporated into the emulation. As neural networks allow for flexible memorization and interpolation, they provide a compact statistical representation of complex data allowing for rapid inferences to be made.

The main alternative to MDN for the emulation of a stochastic model are Gaussian processes [28]. These represent the outputs of a model as a multivariate normal distribution. This allows for the quantification of uncertainty and for covariance between points in the model’s input space. Their underlying assumption of the data being normal can make them restrictive as to the type of data that it can represent, however. We have demonstrated here that an MDN can be applied to both overly-dispersed count data (negative binomial example), as well as bimodal count data with a finite domain (final-size distribution example). These examples would be inappropriate to apply a Gaussian process to and so we have not included a direct comparison here.

The use of an MDN emulator for a stochastic model are two-fold. As the neural network directly learns parameters of the mixture distributions, these may be used directly in the output by for example estimating the mean and variance at each point. The emulator may also use the learned distributions to perform random draws from the emulator representing a realisation of the stochastic process. As the emulator approximates the distribution of the output given model input, this essentially produces a likelihood of the data-point given the model parameters. Such a synthetic likelihood could then conceivably be used in a Bayesian inference scheme, such as in a approximate Bayesian computation [15]. It would be interesting to apply this approach where a model likelihood is computationally intractable.

The training of neural networks can lead to the vanishing / exploding gradient problem [29]. Techniques such as momentum and improved initialisation can help mitigate these issues [30]. Anecdotally, we found that re-initialization with a smaller learning rate generally resolved issues encountered in training. Training of neural networks typically involve a large amount of data. If these can be readily generated from a model then an MDN provides a feasible approach to emulation. However when data is small, either a GP or a simplified neural network may be a more appropriate approach.

When model computation is slow or there is a large number of input parameters, a more efficient sampling scheme of the parameter space may be appropriate [31]. Efficient high-dimensional sampling schemes such as entropy maximization have been implemented previously in GPs [14,32]. As these techniques also involve an approximated likelihood of the data, they could be readily implemented into an MDN scheme, where learning can be conducted in an online fashion.

It is also important to consider the types of appropriate distributions to emulate the model output. Whether they are discrete (e.g. binomial or Poisson) or have finite support (e.g. binomial) can impact the resulting approximation. For example using a mixture of normal distributions to describe a finite population would lead to some probability of the population being negative. When the population is small this would be non-negligible (see supplementary). It is therefore important to understand the nature of the data being approximated, for example a final-size distribution can be well-approximated by a Poisson distribution under certain conditions [33]. Plotting the emulated and real data, either as their summary statistics or as a point cloud as was done here is an important step toward understanding the validity of the emulator approximation.

Neural networks and in particular deep learning has made enormous progress recently, rapidly improving the state-of-the-art in representation of data sets [34]. This has also led to an increase in open software for developing neural networks including Keras and Tensorflow [35,36]. Building from these allow the rapid development of new emulation models and provide the use of established code for the testing and analysis of the trained models. In companion to this article we provide an open access Python library to develop an MDN emulator with example notebooks demonstrating its use. The library also provides details on the exportation of a trained emulator into a web application. For more information, see S2 Appendix.

## Conclusion

Mixture density networks have the potential to be used as emulators for complex epidemiological agent-based and micro-simulation models. These techniques incorporate cutting-edge advances in machine learning that provide the possibility to leverage new software libraries in order to perform fast emulator fitting. Applications can include the building of web interfaces for models as well as in model fitting. We hope this technique will prove useful to the broad epidemiology modelling community and as such have included an accompanying open-source library with examples demonstrating its use.

## Supporting information

### S1 Appendix. Choice of distribution in mixture density network

The mixture density network (MDN) can be fitted using different types of distribution and the choice of distribution is essential in achieving a mixture distribution that matches well with the empirical distribution of the original inputs. In choosing an appropriate distribution, the support of the distribution should be considered such that it matches the original output being used to train the network.

For the final size distribution results (Fig. 3), the final size is given by the total number of individuals infected over the course of the epidemic (calculated as N (∞) – S (∞)). Since, the population size used is *N* = 1000 and the initial conditions are for one infected person, with the rest susceptible, the final size can then take an integer value between 1 and 1000 inclusive.

We test the results for 5 different distributions: normal, gamma, beta, Poisson and binomial. The training data was 10,000 samples of the actual simulated process using input parameters *β* and *γ*, as described in the main manuscript. For the continuous distributions (normal, gamma and beta), in the training process the output final size of the simulation was linearly scaled to be in the interval [0.000001, 0.999999], such that the beta distribution, which has finite support of the interval (0, 1), could replicate these results in the emulation. For the discrete integer distributions (Poisson and binomial), one was subtracted from the simulated output, such that the output training data was between 0 and 999; this means that a final size of 0 cannot be given by the emulated distributions when the data is scaled back to be between 1 and 1000.

For the normal distribution, the emulation has a poor fit, since the re-scaled output can be below 1 or more than 1000 (Fig. S1A). This is exemplified in the two sample Kolmogorov–Smirnov (KS) test (Fig, S1B), where the KS-statistic is much larger than the threshold for the simulated and emulated distributions to be accepted as being the same distribution, for all *R*_0_ values. Additionally, sampling from the emulated distribution and then rounding the results does not significantly improve the quality of the KS test (Fig. S1C).

The gamma distribution shows an improved fit on the normal distribution (Fig. S1D), but since the upper limit is unbounded, the fit in this region remains poor. The model fit is improved further by using the beta distribution, which is bounded at both limits (Fig. S1G). KS tests show that neither distribution are sufficiently similar to the simulated distribution (Figs. S1E and H). However, by rounding after sampling from these distributions and re-performing the KS-test the results appear to be significantly better (Figs. S1F and I). This is because a large proportion of the simulated results will be 1 or 1000, where either the epidemic has died out through stochastic fade-out or the whole population becomes infected and rounding the emulated results achieves this exactly. This effect means the KS-statistic becomes small for small *R*_0_ for the gamma distribution and small for both small and large *R*_0_ for the beta distribution, which is bounded at both limits.

The Poisson distribution, being a discrete distribution achieves a good fit when the final size is small, but for larger final sizes, being unbounded and only specified by one parameter, the fit is much worse (Fig. S1J). This is reflected in the KS test statistic (Fig. S1K).

The binomial distribution provides the closest match to the simulated data since it is both integer valued and bounded at both limits, like the simulated results themselves (Fig. S1L). Indeed, the KS test shows the threshold is met for the simulated and emulated distributions to have come from the same distribution across much of the *R*_0_ value range. Hence, this distribution was used in the main manuscript for the final size distribution.

These results are analogous for other simulated training data – to get the best match in the emulation, the output distribution should be chosen to match the characteristics of the simulated output.

### S2 Appendix

The open-access Python library for the method along with example code and notebooks to reproduce the examples given in this study can be found at the following url: https://github.com/QCaudron/pydra.

## Acknowledgments

The authors gratefully acknowledge funding of the Children’s Investment Fund Foundation. The funder had no role in study design, data collection and analysis, decision to publish, or preparation of the manuscript.

---

**Figure S1:**
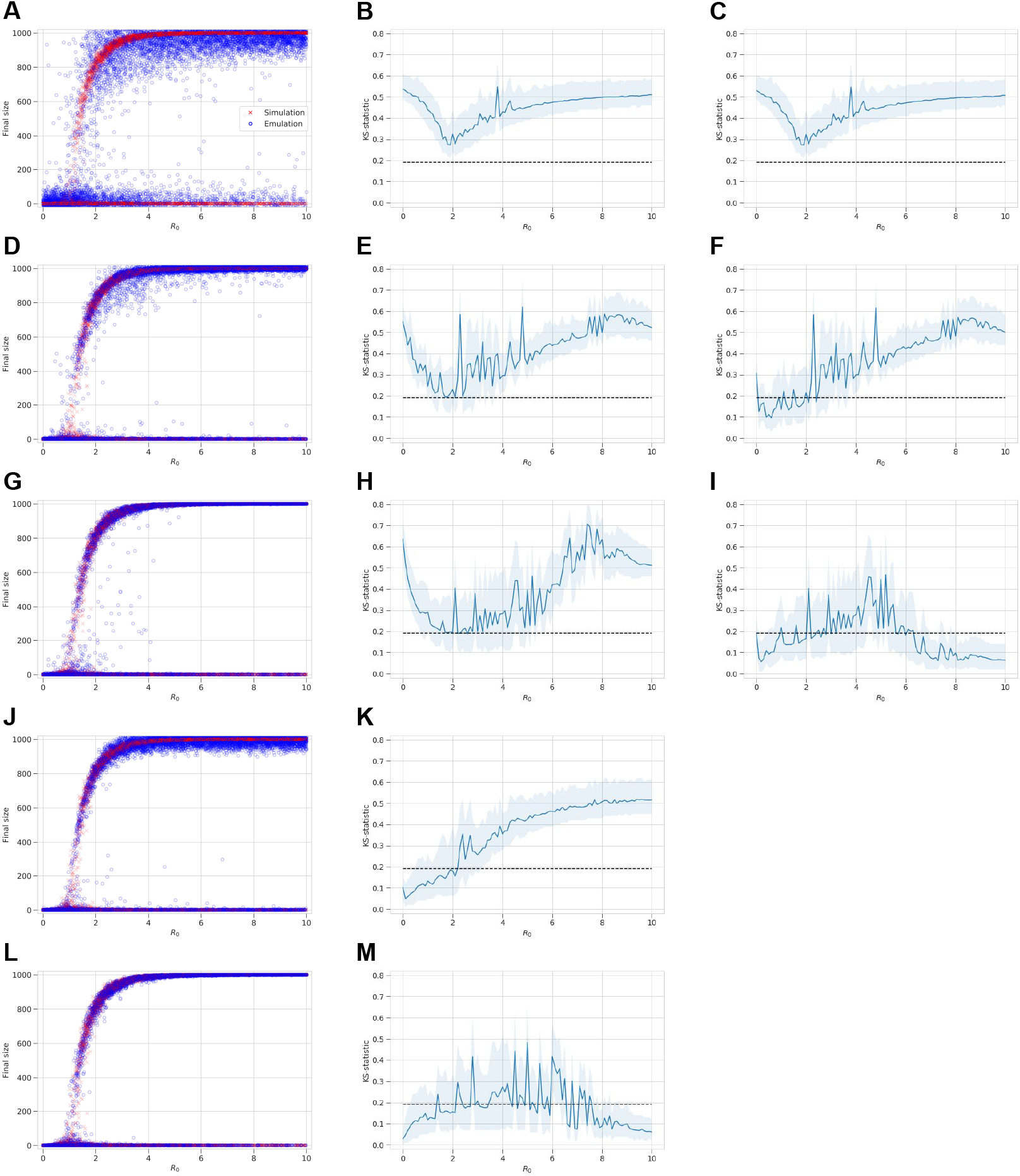
Emulation and simulation comparison for output MDN distributions. **(A-C)** Normal distribution. **(D-F)** Gamma distribution. **(G-I)** Beta distribution. **(J-K)** Poisson distribution. **(L-M)** Binomial distribution. The first column compares the simulated and emulated output empirical distributions, the second column is the results of a two sample Kolmogorov-Smirnov test of the simulated and emulated distributions, and the third column repeats the KS test where the emulated output has been rounded if the chosen output distribution is continuous.

